# Temporal prediction captures retinal spiking responses across animal species

**DOI:** 10.1101/2024.03.26.586771

**Authors:** Luke Taylor, Friedemann Zenke, Andrew J. King, Nicol S. Harper

## Abstract

The retina’s role in visual processing has been viewed as two extremes: an efficient compressor of incoming visual stimuli akin to a camera, or as a predictor of future stimuli. Addressing this dichotomy, we developed a biologically-detailed spiking retinal model trained on natural movies under metabolic-like constraints to either encode the present or to predict future scenes. Our findings reveal that when optimized for efficient prediction **∼**100 ms into the future, the model not only captures retina-like receptive fields and their mosaic-like organizations, but also exhibits complex retinal processes such as latency coding, motion anticipation, differential tuning, and stimulus-omission responses. Notably, the predictive model also more accurately predicts the way retinal ganglion cells respond across different animal species to natural images and movies. Our findings demonstrate that the retina is not merely a compressor of visual input, but rather is fundamentally organized to provide the brain with foresight into the visual world.

## Introduction

Retinal ganglion cells (RGCs) respond to different patterns of light [1, 2] and send a representation of the visual world to the brain [3], spatially downsampled by about 100 times from the photoreceptors. It remains a open question what information this pattern of activity represents. One hypothesis posits that the retina efficiently compresses incoming light into a downsampled neural code [4–10] - like a camera capturing incoming light. Another hypothesis is that the retina efficiently signals features within sensory input that are predictive of the future [11–15] - like a crystal ball foretelling the future. The concept of prediction is thought to be fundamental to many aspects of sensory processing, where the predominant focus has been on the role of cortical circuits in predicting sensory inputs [16, 17].

Normative modeling is a commonly used approach for exploring such coding questions [18, 19]. This involves optimizing a model (in our case of the retina) for a particular goal, for example efficient compression or prediction, and comparing the resulting phenomena within the model to the biology. Previous normative models of the retina have mostly explored the efficient compression of natural images by employing model units that output firing rates [4–8]. These models have successfully captured the different types of retina-like spatial receptive fields (RFs) and their mosaic-like organizations. However, they lack particular biological features, such as the spiking responses of RGCs and their recurrent interactions [20]. Modeling spikes is important since RGCs may encode visual stimuli using the relative timing of spikes, which therefore play an important role in sensory transmission [21]. Furthermore, these models have not been shown to capture more complex retinal processes, such as stimulus-omission [22, 23] and motion-anticipation [24] responses or differential motion tuning [25]. Most importantly, and pivotal to the compression versus prediction dichotomy, it remains unclear to what extent a spiking model optimized for prediction compares to the retina.

To address these shortcomings, we developed a spiking retinal model, trained on natural movies under metabolic-like constraints, to either encode the present or to predict the future, from recent spike activity. Like prior non-spiking normative investigations [4–8, 15], we found that our spiking model optimized for prediction was able to capture retina-like RFs and their spatial organizations. Extending beyond these results, we found that our model also exhibited retina-like latency coding [21] and spike statistics in response to natural images and movies. Furthermore, the predictive model captured complex retinal processes of motion tuning and sensitivity to the unexpected absence of a stimulus (stimulus-omission responses) [22, 23]. In contrast, the model optimized for efficient compression did not account for complex retinal processes, like stimulus-omission responses. Notably, we found the model optimized for prediction ∼ 100 ms into the future to more accurately predict RGC responses across different animal species to both natural images and movies. Our findings therefore suggest that the retina in mammals and amphibians has evolved to supply the brain with temporally-predictive features about the visual world.

## Results

### The spiking retinal model

The retina consists of three cellular layers: the photoreceptors (which transduce light), the bipolar cells (an intermediate cell layer), and the RGCs (the spiking output layer), interspersed by the horizontal and amacrine cell inhibitory interneurons [20] (Fig. 1a). We modelled this circuit using a single hidden layer of recurrently-connected spiking units (Fig. 1b), where the spiking units are equivalent to the RGCs. All spiking units were modelled using the leaky integrate-and-fire model [26], which captures the key biophysical nature of real neurons: integrate incoming current, and fire a spike when the firing threshold is reached (Fig. 1c). Pre-ganglionic cells (the photoreceptors, horizontal and bipolar cells) appear to mostly respond to incoming light in a linear fashion [27]. We thus opted to model their transformation using a linear mapping, whose output was fed as input current to the RGC-like spiking units. Similar to prior modelling work [28], we included recurrent connectivity to capture the interactions between RGC types, arising from electrical synapses between RGCs [29, 30] and electrical and chemical synaptic connections between RGCs via amacrine cells [20, 29]. We also noise-perturbed the input stimulus and each unit’s membrane potential with Gaussian noise, to capture the noise characteristics of the photoreceptors and RGCs [31, 32]. We chose these noise sources so that the model’s spike train variability matched that of the RGCs across repeated stimulus presentations. As in the retina, the model’s spike trains were almost identical in response to uniform full-field illumination with randomly varying intensity over multiple repeats [3, 33], and the model’s response variability was sub-Poisson with Fano factors mostly below unity [33–35] (Extended Data Fig. 1).

**Fig. 1:**
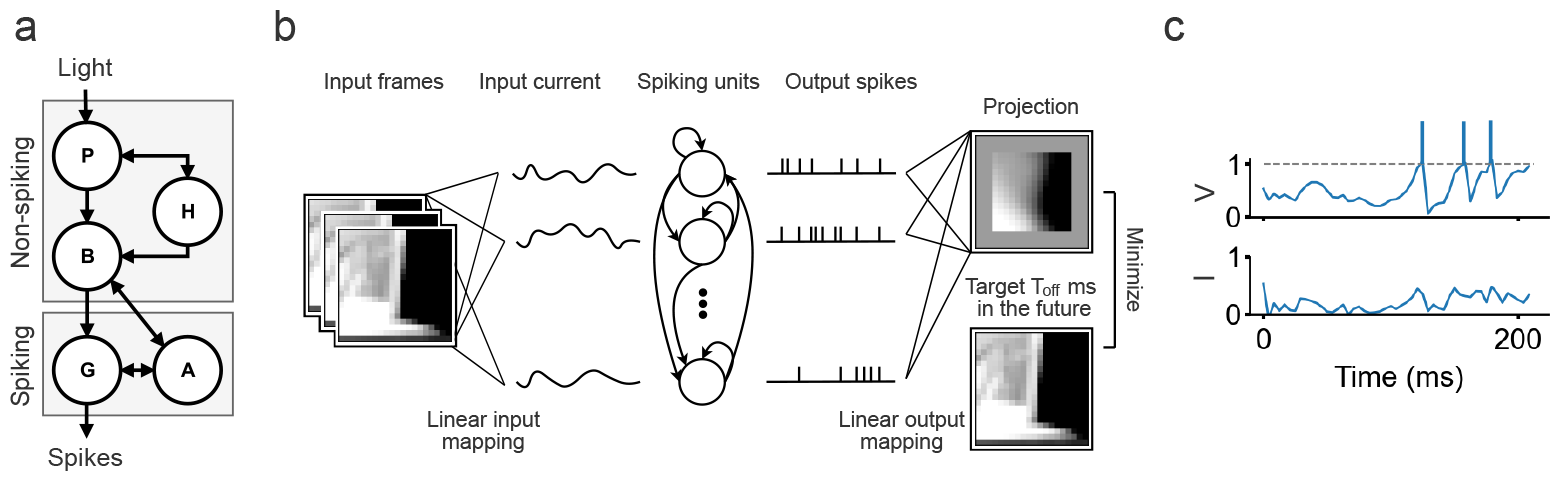
The spiking retinal model. **a**. Schematic of the retinal circuit with its different cell types (P=Photoreceptor, H=Horizontal cell, B=Bipolar cell, A=Amacrine cell, G=Ganglion cell) and respective connections, with the arrow heads indicating the flow of visual information. **b**. Schematic of the retinal model. The model takes a sequence of movie frames as input and linearly transforms them into an input current that is fed into a single hidden layer of recurrently-connected spiking LIF units. Photoreceptor-like noise was added to the input image and ganglion-cell-like noise was added to the input current. The spike output of these units was used to linearly construct a projection, either of the present or the future, of the input stimulus. The linear transformation approximates the transformation of the pre-ganglionic cells in the top gray box in **a**., and the recurrently-connected spiking units model the ganglion cells, and their interactions with the amacrine cells, in the bottom gray box. **c**. Example activity traces of one of the LIF units over time: input current *I* charges the membrane potential *V* and elicits a spike if the firing threshold (dotted line) is crossed.

The model was trained on diverse natural movies, with the objective of estimating a spatial frame *T*_off_ ms into the future, or at the present, using a linear readout from past spike activity. We explored different prediction offset values *T*_off_ to examine whether an encoding (*T*_off_ = 0 ms) or predictive (*T*_off_ *>* 0 ms) objective better captures the sensory processing of the retina. Lastly, we regularized the model’s connectivity during training using a metabolic-like cost function, which penalized individual synaptic-like connections by their cost in energy, approximated as the absolute value of a connection weighted by its incoming activity over time (see Methods). Unless otherwise mentioned, we report our results using the model optimized for prediction of *T*_off_ = 128ms into the future.

### Retina-like spike statistics to natural images and movies

First, we checked that our model could produce retina-like spike trains. We compared the spiking activity of the model units to RGC activity across different animal species in response to natural images and movies. Here, we used publicly available recordings from the retina in macaque, marmoset, mouse and salamander [35–37]. We found our model to exhibit similar spike activity to the RGCs across each of these species. The retina qualitatively exhibits more sparse and regular firing activity in response to natural images [35] (Fig. 2a) compared to natural movies [37] (Fig. 2b). We found that our model exhibits similar firing patterns for the different types of input.

**Fig. 2:**
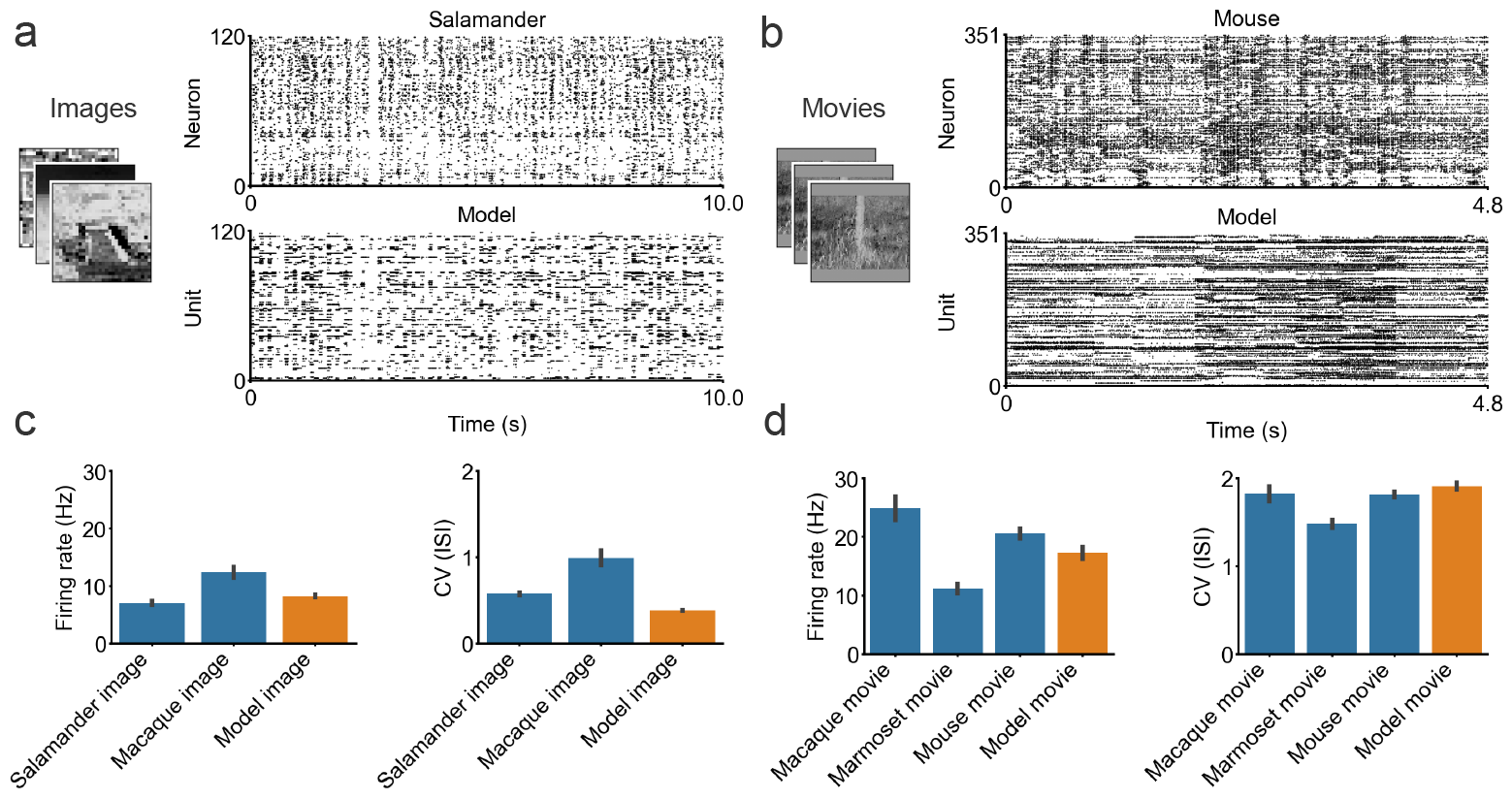
Spike responses to natural images and movies. **a**. Raster plot from the salamander retina (top) and the model (bottom) showing responses to the same natural images (left). **b**. Raster plot from the mouse retina (top) and the model (bottom) to the same natural movies (left). **c**. Left: Mean firing rate of RGCs from different species (blue) and model units (orange) evoked by natural images. Right: Corresponding mean coefficient of variation of the interspike intervals CV(ISI) of the neurons and model units to the natural images. **d**. Same as **c**. but with the statistics instead calculated using the responses to natural movies. The salamander data were obtained from [35]; the macaque image and movie data were obtained from [36], and the marmoset and mouse movie data were obtained from [37]. The number of model units was randomly sampled for plots **a**. and **b**. to match the number of neurons. All bars plot the mean and s.e.m.

We quantified the sparsity and variability in the spike responses to the natural images and movies across the different animal species and the model. We measured the spike response sparsity and variability by respectively calculating the mean firing rate and the mean coefficient of variation of the interspike interval CV(ISI). The CV(ISI) measures the variability in a spike train, where *CV* (*ISI*) *<* 1 corresponds to a more regular firing pattern than a Poission process and *CV* (*ISI*) *>* 1 a less regular one [26] (see Methods). Both the mean firing rate and CV(ISI) are lower across the animal species in response to the natural images (Fig. 2c) than to the natural movies (Fig. 2d). Similarly, we found the model units to exhibit a lower firing rate (images 8.27Hz vs movies 17.27Hz; *p* = 6.01×10^*−*14^, one-sided Mann-Whitney U test) and CV(ISI) (images 0.38 vs movies 1.91; *p* = 1.87×10^*−*166^, one-sided Mann-Whitney U test) to the images compared to the movies. We used the same image dataset from the salamander recordings [35] to quantify the model’s response statistics to natural images, and used our held-out movie test set to calculate the model’s response statistics to natural movies.

### Emergence of major retina-like cells

We analyzed the functional type of each model unit, as one would categorize the functional types of RGCs [35], by mapping the spatiotemporal receptive fields (RFs) of the model units using a spike-triggered average to white-noise. We fitted a 2D Gaussian function to each unit’s spatial RF and projected the spatial RF size and dimensionality-reduced RF temporal profile onto a 2D scatter plot. We then separated the units into functional groups using k-means clustering (see Methods).

We found the model units to separate into four functional classes (Fig. 3a), each resembling one of the major cell types found in the primate retina, which together constitute over 95% of the RGCs in the central retina [38]: ON parasol and midget cells and their OFF type counterparts [38, 39] (Fig. 3b). Just like the real retina, the parasol-like units had a larger spatial structure and a more biphasic temporal profile than the midget-like units [40, 41]. We also found the model parasol- and midget-like units to behave similarly to their biological counterparts in response to flashes of light (Fig. 3c): the ON and OFF parasol-like units responded with a transient burst of spikesto changes in light intensity, and the ON and OFF midget-like units responded with a sustained train of spikes (with the ON-type cells responding to the light onset, and the OFF-type cells responding to the light offset) [42, 43]. As reported in the retina across different species [44–46], the RFs of the different unit types in the model tiled visual space, forming mosaic-like organizations (Fig. 3d). We also found fewer parasol-than midget-like units, and fewer ON-than OFF-type units within the model (Fig. 3e). This aligns with experimental reports of midget cells being more plentiful than parasol cells [38, 47], and OFF-type cells occurring more frequently than ON-type cells [41, 48]. Lastly, the model units exhibit a heterogeneous membrane time constant distribution on the scale of tens of milliseconds, which has similarly been reported for cat RGCs [49] (Fig. 3f).

**Fig. 3:**
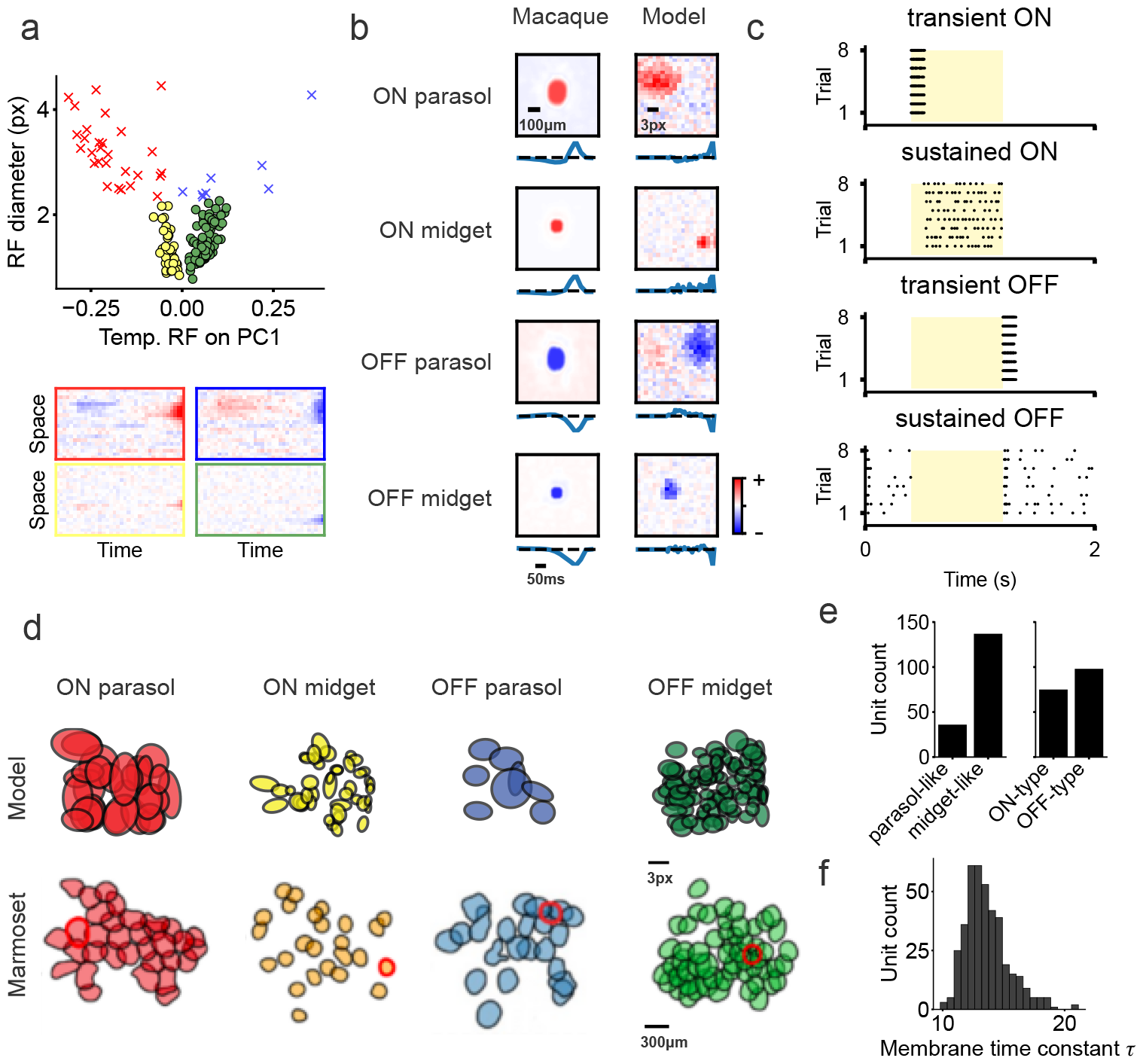
Retina-like RFs and their mosaic organization. **a**. Scatter plot of model unit RF diameter (obtained from a 2D Gaussian fit) vs the first principal component of the dimensionality-reduced RF temporal profile, with the spatiotemporal RFs obtained from a spike-triggered average to white-noise. Separate colors and shapes denote distinct functional groups (obtained using k-means clustering), with red and blue crosses denoting ON-type and OFF-type parasol-like units respectively and yellow and green dots denoting ON-type and OFF-type midget-like units respectively. Example 2D spatiotemporal RFs from each group are plotted below: left are ON-type units, right are OFF-type units, top are parasol-like units and bottom are midget-like units. **b**. Example spatial RFs, with corresponding temporal profile plotted below, from the macaque retina [4] (left) and the model (right), where red and blue colors denote high and low power respectively. **c**. Light-flash responses of the model units depicted in **b**. (yellow rectangle denotes the duration of the flash of light). **d**. RF mosaics of the different model unit types (top) with the corresponding RF mosaics of the different marmoset RGC types [37] (bottom). **e**. Left: number of model units that are parasol- and midget-like. Right: number of units which are ON- and OFF-type. **f**. Membrane time constant distribution over all model units.

### Texture, differential and anticipatory motion selectivity

We quantified the tuning characteristics of the model units and found that they exhibited several well-known motion-tuning characteristics of RGCs. Some RGCs in different species have been reported to be selectively tuned to the orientation and direction of moving objects, with a preference for motion along the cardinal axes [50–57]. Similar to the biology, we found some units to be selective for orientation (16.37% of units with *OSI >* 0.5) and some selective for direction (3.48% of units with *DSI >* 0.5), with a preference for motion along the horizontal and vertical axes (Fig. 4a). A well-known class of RGCs, known as Y-type cells (∼ 3% of ganglion cells in cats [58]), are tuned to the general motion of textures of high spatial frequency, but not their static presentation [20, 59–61]. Similar to these neurons, we found units in the model (7.66% of units) that did not respond to a stationary grating of high spatial frequency (for each unit’s preferred orientation), yet fired as soon as the grating moved (Fig. 4b). Some RGCs, known as object-motion-sensitive (OMS) cells, exhibit differential motion selectivity, where a neuron remains silent when an image (or grating) moves across the retina, yet fires if the motion in the RF center differs from that in the wider surround (*e*.*g*. by masking out the wider surround) [25, 52, 62, 63]. We also found such OMS tuning in some of the model units (Fig. 4c). Certain RGCs have an anticipation-like response to moving objects (like a bar moving left or right), where the neuron fires even before the object crosses the neuron’s RF, whereas there is a delayed response to simply flashing the bar onto the RF [24]. Similarly, we found certain units in our model to also have an anticipation-like response to a moving bar (Fig. 4d).

**Fig. 4:**
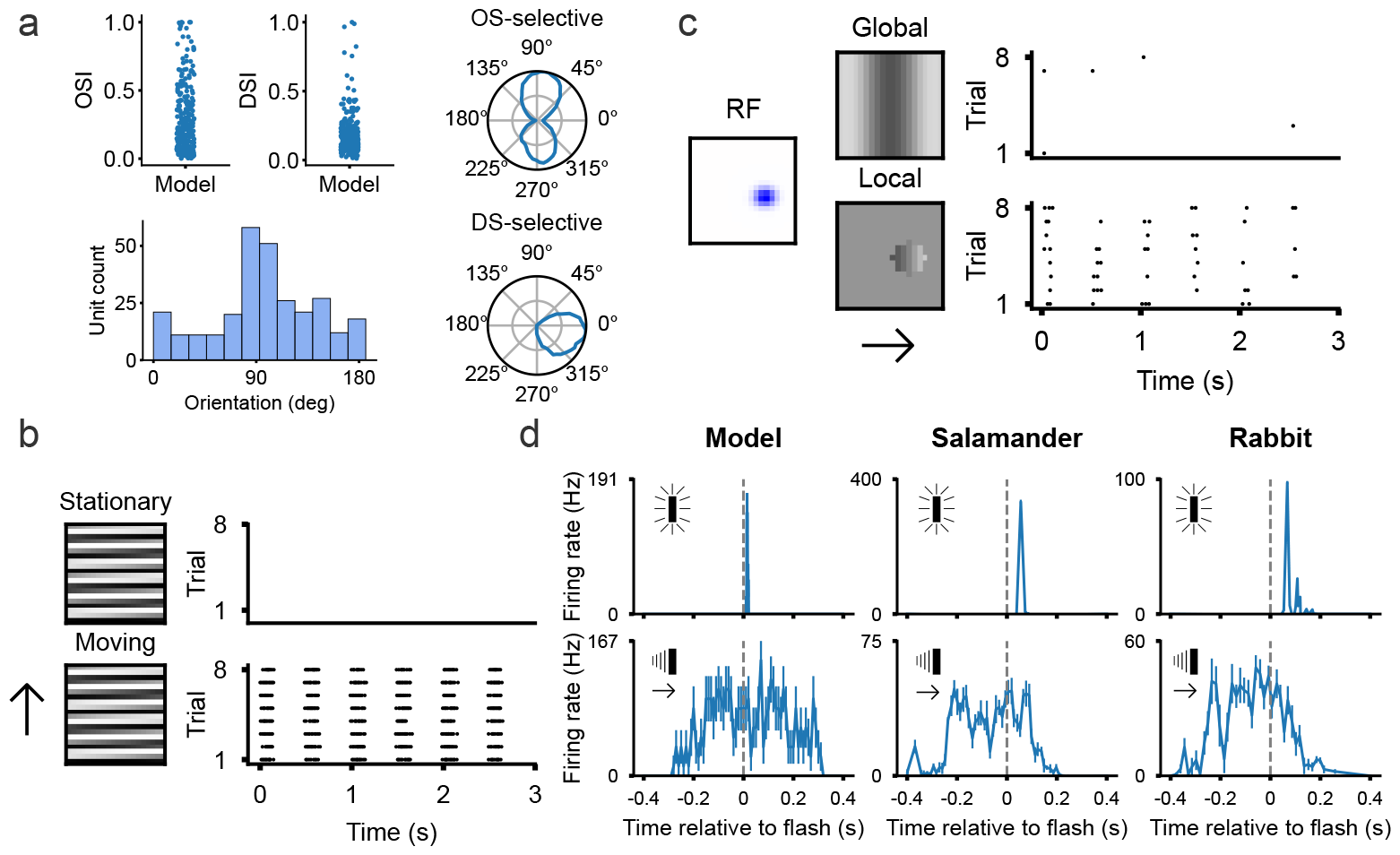
Motion tuning. **a.** Top left: model unit orientation- and direction-selectivity distributions (values closer to unity denote stronger tuning). Bottom left: distribution of orientation tuning for the model units. Right: sample tuning curves of an orientation-selective and direction-selective unit. **b**. Spike raster of a model motion-sensitive unit, with no response to a stationary grating of high spatial frequency (top), and a response to the grating moving upwards (bottom). **c**. Spike raster of an object-motion-sensitive model unit, with no clear response to the grating moving over the entire visual space (top), and a stronger response to the same spatially localized grating moving within the model unit’s RF (bottom). **d**. Top: responses of a single model unit, and a single salamander and rabbit retinal ganglion cell to a briefly flashed bar (∼ 15 ms) within their RFs. Bottom: Anticipation-like responses in the same model unit and RGCs to a rightward moving bar (∼ 0.44 mm s^*−*1^ - see Methods) before the bar crosses the RFs. The bars were aligned at time zero and error bars denote the s.e. over repeated stimulus presentations, with the salamander and rabbit data adapted from [24].

### Emergence of a relative spike latency code and stimulus omission responses

A contentious and open question in neuroscience is how neurons transmit information using their spikes [64]. This has historically been cast as a code of two extremes, where neurons either communicate using the number of spikes evoked (*i*.*e*. a rate code) or the timing of spikes (*i*.*e*. a temporal code). These strategies are not necessarily mutually exclusive [65], and different neurons likely employ different coding strategies depending on their location within the nervous system. In the retina, there is strong evidence that certain RGCs employ a temporal-like code, encoding the onset of a new visual scene using the relative timing of the first spikes. Reconstructing natural images from such a differential spike latency of fast OFF RGCs, rather than their spike counts, qualitatively results in a clearer image reconstruction [21]. We examined these coding strategies in our model and similarly found a rate-based reconstruction to qualitatively be more blurry, noisy and with erroneously overemphasized edges than a latency-based reconstruction (Fig. 5a). A differential spike latency code can transmit new visual information using very few spikes, which is both energy efficient and fast. This is particularly important, as our gaze only remains fixed for a fraction of a second, before rapid saccadic eye movements induce a new visual scene onto the retina [66, 67].

**Fig. 5:**
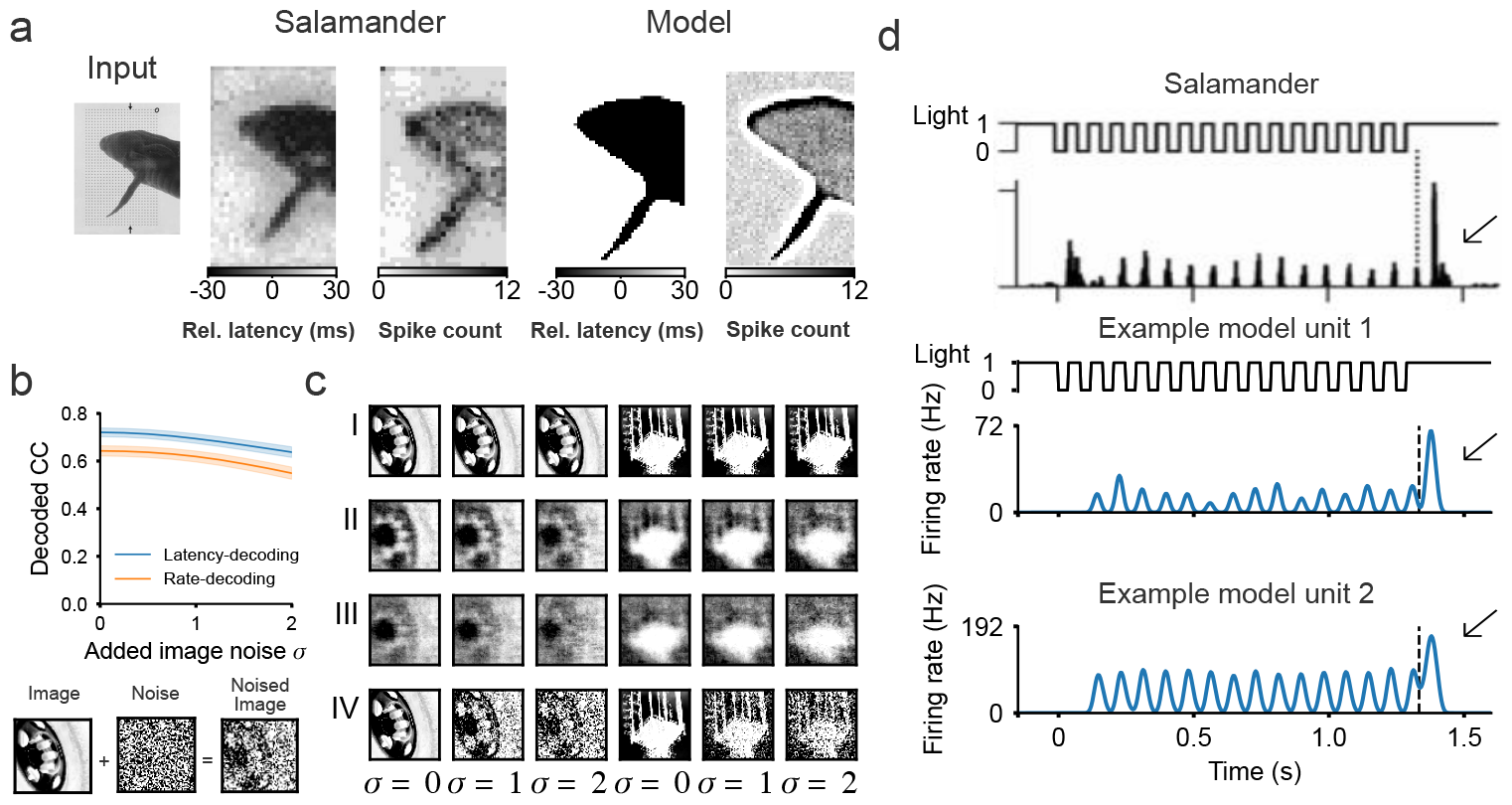
Latency and stimulus-omission responses. **a**. Reconstruction of an input image using the relative spike latencies and the spike counts from a single OFF-type neuron from the salamander retina [21] and from a single OFF-type model unit. **b**. Top: mean decoding accuracy (Pearson correlation coefficient) in reconstructing natural images from noise-corrupted inputs using a linear transformation of the population model activity with a latency code (blue line) and a rate code (orange line) (the shaded area is s.e.). Bottom: illustration of noise-corrupting the input images. **c**. Example image reconstructions from **b**., I: target image, II: latency-decoding, III: rate-decoding, IV: noised-image. **d**. Firing rate of a single salamander RGC [22] (top) and of two example model units (bottom) during a sequence of 16 40ms-flashes presented at 12Hz. The dashed line marks the time when the flash sequence was halted, with the arrow indicating the omitted-stimulus response. Omission stimulus responses were also present at different flash rates, as well as when flashes were omitted in the middle of the sequence (Extended Data Fig. 2).

Most of our understanding of the retina’s neural code is at a single-cell level based on responses to artificial stimuli, like drifting gratings [20, 68]. It is less well understood how natural stimuli are encoded by the population activity. Additionally, it remains to be understood how the different coding strategies compare to each other in the presence of varying levels of noise. This is important, as the retina likely employs a noise-robust coding strategy, given the ubiquity of noise in the retina and the visual environment [69]. To explore this, we tested how well we could reconstruct natural images, by inputting their noise-corrupted counterparts into the model and linearly decoding the images from the resulting rate and latency codes of the population model spike activity. We found that the latency-decoding strategy more robustly decoded the noise-corrupted natural images than the rate-decoding strategy (Fig. 5b), qualitatively producing clearer image reconstructions, even under high levels of noise corruption (Fig. 5c).

Certain RGCs in the salamander and mouse retina have been reported to produce an omitted-stimulus response (OSR), generally thought to encode prediction or prediction errors [70]. RGCs exhibiting OSRs fire strongly to a violation in a periodic temporal pattern, such as an omitted flash within a steady dark flashing sequence [22]. Such neurons either respond weakly (20% of neurons) or not at all (22% of neurons) during the flash sequence, as reported in the salamander retina [23]. We found that some of the model units (7.66%) produced an OSR to a violation in a periodic sequence flashing at a rate of 12 Hz (see Methods). These units’ responses were sustained throughout the flash sequence, and greatly increased at the end of the sequence, around the expected time of the response to the omitted flash (Fig. 5d). We also found OSRs to be present for slower (8 Hz) and faster (16 Hz) flash sequences, as well as when flashes were omitted in the middle of the sequence (Extended Data Fig. 2). Interestingly, OSR responses were only present in the model optimized for temporal prediction and not in the model optimized for compression.

### A predictive retinal model captures neural responses across animal species

We assessed whether optimizing the retinal model to encode the present or the future better predicts the activity of RGCs across different animal species in response to natural images and movies. We used five publicly available datasets of RGC recordings in macaque, marmoset, mouse, and salamander [35–37]. For every dataset and retinal model, we fitted a linear-nonlinear readout from the retinal model to predict the neural responses [71] and quantified the performance of each neuron’s fit as the normalized correlation coefficient (*CC*_*norm*_) between the predicted and the target neuron response on a held-out validation set [72]. As commonly done [73], we performed the fits from a dimensionality-reduced latent representation of the model’s hidden-unit activity in response to the different dataset inputs (Fig. 6a; see Methods).

**Fig. 6:**
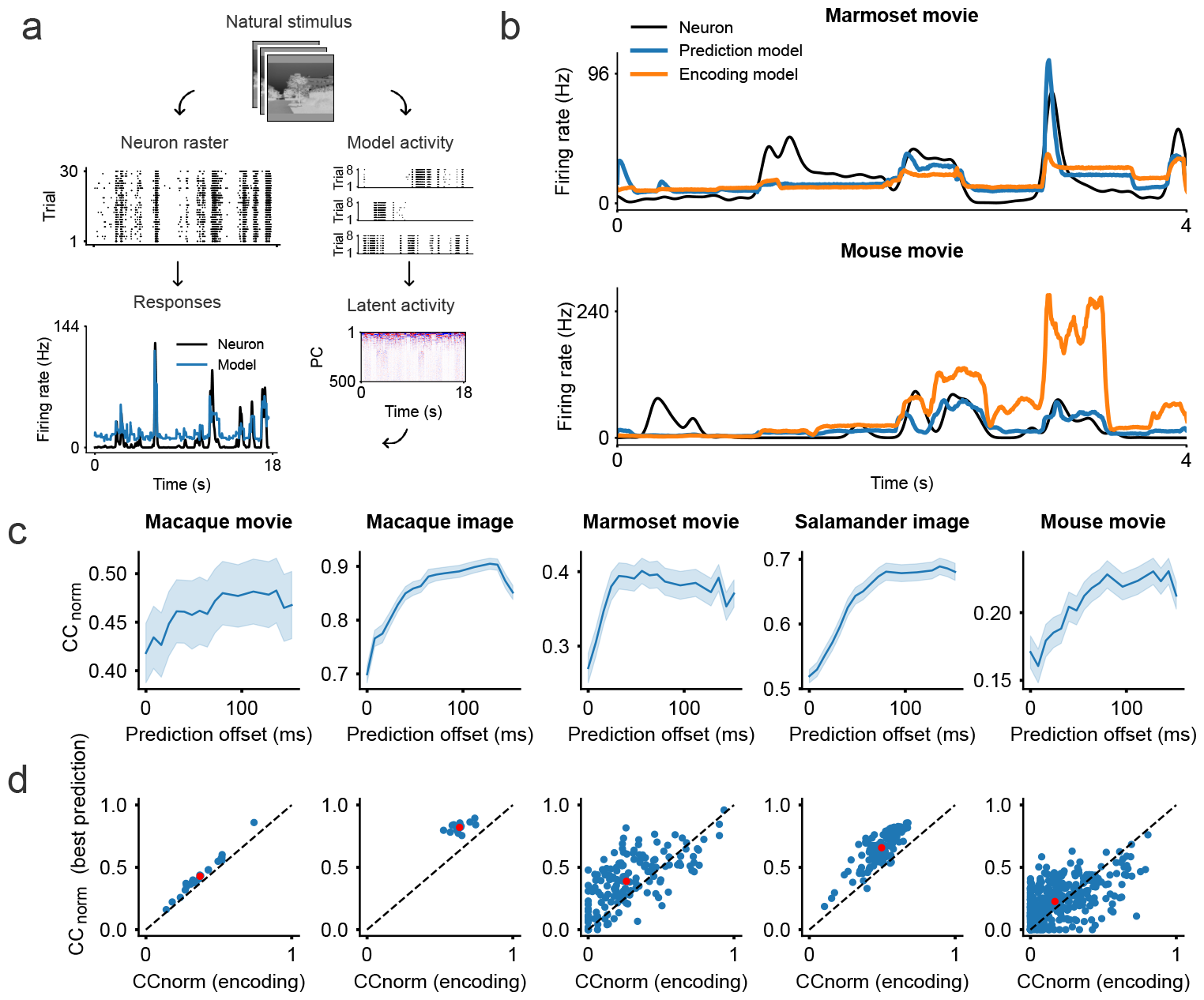
Predicting RGC responses to natural images and movies using the spiking models. **a**. Each retinal model (trained to encode the present or predict different offsets into the future) was fitted to predict the neural responses across multiple natural stimuli in different retinal datasets. Model unit activity was obtained for each stimulus set used in these studies and downsampled into a latent activity space using principal component analyses, from which the neural responses were predicted using a regularized linear-nonlinear readout (see Methods). All results report the normalized correlation coefficient between the predicted and the target neural responses, computed on a held-out validation set. **b**. Sample neural response (black line) of a single neuron in the marmoset retina (top) and mouse retina (bottom) to natural movie stimuli. Predicted responses of the predictive model (blue line) and encoding model (orange line) are superimposed. **c**. Neural prediction scores across the different retinal datasets and retinal models, as a function of the prediction offset (graphs plot the mean and s.e. over neurons). **d**. Neuron prediction scores of the best predictive model (y-axis, optimal offset) against the encoding model (x-axis, 0 ms offset) across the different datasets (with the red dot plotting the centroid).

Across all animal species and stimulus types, we found the retinal model optimized for prediction to more accurately predict the neural responses. The predictive model appeared to qualitatively match the neural responses, particularly at the peak firing rates (Fig. 6b). Notably, the mean *CC*_*norm*_ steadily increased across all datasets for increasing model prediction offset (*i*.*e*. the further the retinal model predicted into the future, the better the neural fits), up until a maximum score was reached, after which prediction performance declined (Fig. 6c). Across the datasets, the maximum *CC*_*norm*_ was obtained using the retinal model optimized to predict 56 − 128ms into the future. Lastly, the higher *CC*_*norm*_ for prediction was driven by the majority of neurons - and not some small subset (Fig. 6d).

## Discussion

We set out to uncover the principles underlying the sensory transformation performed by the retina, from the incoming light to the outgoing spikes conducted by the optic nerve. Does the retina efficiently compress incoming input [4–8] or efficiently extract sensory features predictive of the future [11–15]? To examine this, we implemented a biologically-detailed spiking model of the retina, which we separately optimized for efficient compression or efficient prediction using natural movie stimuli. We found the model optimized for prediction to exhibit various retina-like phenomena, not all of which were captured by the model optimized for efficient compression. Notably, the predictive model more accurately predicted RGC responses to both natural images and movies across different animal species. These findings suggest an evolutionary conserved role of the retina in informing the brain about temporally-predictive features in the visual world.

Normative approaches enable the underlying principles behind the sensory transformations of a biological circuit to be identified [18, 19]. Normative models of the retina have been constructed and optimized for goals such as compression or prediction. Prior studies developed models of the retina using non-spiking units optimized for efficient coding, by maximizing the mutual information between the sensory input and model unit firing rates [5–8]. These models have accounted for the retina’s spatial RFs [4–8], their temporal profiles [4, 6], and their mosaic-like organization [5–8].

We asked whether normative principles such as compression or prediction explain retinal properties when spikes are used for communication. To this end we developed a retinal model with units that output individual spikes, providing greater biological realism than prior work that used firing rate outputs [4–8]. This is particularly important, as the retina more likely encodes sensory input using spike times than firing rates [21]. Our model also includes recurrent connectivity to capture the interplay known to exist between ganglion cells, whereas other models omit recurrence [4–8]. Like prior studies [5–8], our model incorporates input and output noise to model the noisy phototransduction and the noisy spike generation of RGCs. We trained our model on natural movie recordings, whereas previous work has omitted temporal dynamics by training on natural images [5, 7, 8] (although see [6]). Our model was also trained to minimize a detailed metabolic-like loss, approximating synaptic energy requirements, unlike previous studies, which have simply constrained model firing rates [4–8].

We explored the longstanding hypothesis of the retina performing prediction, in contrast to studies focusing exclusively on the retina encoding the present visual scene [5–8]. Like prior work, our model captures the spatial RFs, their temporal dynamics, and mosaic-like spatial organization. Unique to our work, we contrasted the spiking dynamics of our model to the spiking dynamics of RGCs. We found our model to exhibit retina-like spike statistics to natural images and movies, as quantified using firing rates and measures of spike variability. Notably, our model more accurately encoded sensory input using the relative spike latencies rather than firing rates, as has also been reported in the retina [21].

Our model exhibited several retinal phenomena unaccounted for by other normative studies of the retina. This includes finding certain units tuned to the direction and orientation of moving gratings, as has similarly been reported in the retina [74]; units that exclusively fire to high-spatial-frequency gratings only if they move, like Y-type cells in the retina [20]; units tuned for the differential motion of objects versus their background [25]; units tuned for the anticipation of moving objects [24]; and units tuned for stimulus omission within a temporal sequence of flashing lights [22, 23]. Certain complex retinal phenomena (like omission and anticipation responses) have been shown to emerge in non-spiking encoding models of the retina [75, 76]. These phenomena emerge as a consequence of fitting models to retinal responses, whereas our normative approach rather examines whether these phenomena can be explained as a consequence of underlying principles like compression or prediction.

Salisbury and Palmer [12] hypothesized that certain nonlinear retinal processes may be connected to prediction. Supporting this view, we found the stimulus-omission responses to emerge only when our model was optimized for efficient prediction and not for efficient compression of the present. Lastly, we addressed the challenging problem of predicting RGC responses to natural images and movies across different animal species. We found that the model optimized for efficient sensory prediction more accurately predicted neural responses than the model optimized for efficient sensory encoding of the present. Interestingly, the best performing model was trained to predict on a timescale of ∼ 50−100ms, similar to the latency of the photoreceptor responses [77, 78]. Furthermore, this duration also corresponds to the duration at which retinal spikes contain the most information about the sensory future, as estimated in a prior study [11].

While the use of spikes and recurrent connections in our model increased realism over prior models, there remains scope to incorporate more of the retina’s biological complexities and processes. RGCs are known to adapt to stimulus contrast [27, 79, 80], which could potentially be modelled using adaptive leaky integrate-and-fire neurons [26, 81]. We could also include constraints on the recurrent connectivity, such as implementing the inhibitory recurrent connections via explicitly modeled amacrine cells [20, 29] and other recurrency via gap-junctions between the spiking units to better emulate the electrical coupling between the RGCs [29, 30]. Furthermore, we modeled the photoreceptors and bipolar cells as a linear transformation, given reports of their near linear integration properties [27]. However, bipolar cells can exhibit some non-linear integration [82], thus modeling them with an additional hidden layer of graded units could be considered.

We trained our model on randomly sampled gray-scale normalized movie patches. We could instead train our model using positive-only pixel values to capture the positive-only nature of light intensity, or with color movies, to examine the retina’s processing of color [83]. Future extensions could also develop and train the model on the entire visual scene rather than patches. This could be a convolutional model [84], or better yet, a model composed of overlapping unique patches, each operating at a different spatial location within the visual frame. This patch model may capture how the tuning properties of RGCs differ across spatial positions, such in their direction-selectivity [85]. However, training such large spiking models remains a challenging problem due to speed and memory constraints (although see [81, 86]).

We could also explore the effect of noise in the model. The RFs in our model lack a clear surround, whereas the RFs of RGCs have typically been characterized as center-surround [20]. Perhaps learning the noise sources in the model, instead of hardcoding them, would result in RFs with surrounds [7]. However, it remains unclear to what degree RGCs exhibit RF surrounds, as several studies report RFs with little to no surrounds [41, 87]. It is possible that these differences result from the way the RFs are mapped, where spike-triggered averages -as employed in our study - have been shown to result in RFs with less defined surrounds [88].

Our model might have clinical implications in the development of retinal prosthetics. Current prosthetics use standard computer vision algorithms to translate camera signals into a stimulatory code [89, 90], which fall short of delivering a normal visual experience [91]. We anticipate that the sensory transformation implemented in our model could aid in delivering a more RGC-like stimulatory signal.

In conclusion, consistent with the capacity of temporal prediction models to explain phenomena in visual [15, 92] and auditory cortex [92], this study adds to the evidence that sensory systems are geared towards prediction of future input. Optimization of sensory systems for temporal prediction enables estimation of the world in the future when action will occur, compensating for transduction, processing and motor delays, while also eliminating irrelevant information and extracting underlying variables [17, 92]. Thus, temporal prediction may be a governing principle of sensory systems.

## Methods

### Spiking retinal model

#### Training data

We recorded a natural visual stimulus dataset for model training. Various types of scenes were obtained, categorized as *flow* (camera moving through space), *pan* (camera panning through space), *still* (fixed camera recording a still scene), and *still moving* (fixed camera recording an active scene). A total of 39 clips of 9.6 second duration were recorded at a frame rate of 120Hz and a spatial extent of 720×1280px. The dataset was divided into a training and test set, both containing a balanced number of the different recording types (training set: 12 *flow*, 8 *pan*, 3 *still* and 8 *still moving* clips; test set: 3 *flow*, 2 *pan*, 1 *still* and 2 *still moving* clips). All clips were pre-processed by converting the RGB channels to grayscale (using a weighted average of the color channels of R:G:B = 30:59:11 [35]). They were then bilinearly downsampled to spatial dimensions of 144×256 and centrally cropped to 140 ×240. The clips were then normalized by subtracting the mean and dividing by the standard deviation of the training set pixels. Finally, the temporal resolution was artificially increased to 240Hz by duplicating every frame. The model was trained on randomly sampled 20×20 pixel spatial patches of temporally non-overlapping subsamples of 320ms in duration (*i*.*e*. 77 frames). We artificially doubled the training data using data augmentation, by flipping each clip around the vertical axis [93].

#### Spiking neurons

We modelled the RGCs in the retina using a single hidden layer of *N* = 400 recurrently-connected spiking units, comprised of a series of computational steps:

Step 1, photoreceptor noise: to emulate the noisiness of the photoreceptors [31], we added Gaussian-sampled noise *ϵ*_*p*_ ∼ 𝒩 (0, 0.01) to every pixel within the input movie stimulus *x* ∈ ^*T ×H×W*^ (for *T* = 77 simulation time steps of *H* = 20 pixels and *W* = 20 pixels of spatial height and width, respectively), to produce noisy input movie stimulus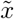 .

Step 2, input current: the input current *I*_*i*_[*t*] ∈ ℝ to unit *i* at time *t* is obtained from the noisy-input movie stimulus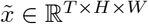, the model output spikes *S*[*t* − 1] ∈ ℝ^*N*^ and a bias term *b*_*i*_ ∈ ℝ.

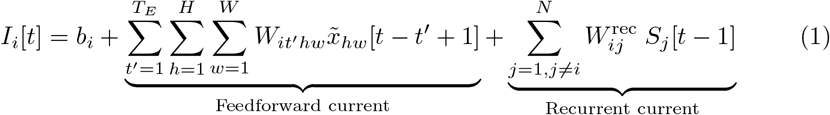

The feedforward connectivity 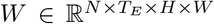 linearly maps the noisy-input movie stimulus to a feedforward current contribution and captures the transformation of the pre-RGC cells. We set the span of latencies to *T*_*E*_ = 30 time steps (125ms) in the feedforward connectivity, consistent with the potentially wide integration times of the photoreceptors [78]. The recurrent connectivity *W*^rec^ ∈ ℝ^*N ×N*^ (autapse connections were excluded) maps the output spikes into a recurrent current contribution to capture the ganglion-to-ganglion cell and ganglion-to-amacrine-to-ganglion cell interactions. Most of the connections between the amacrine cells and RGCs are synaptic [20], with the majority of the amacrine cells using spikes, rather than graded potentials for communication [94, 95], hence the modelling decision to construct the recurrent connectivity from the spiking unit output, rather than the membrane potential via gap-like junctions.

Step 3, RGC noise: to emulate RGC noise [32], we multiplicatively perturbed the input current *I*_*i*_[*t*] of unit *i* at every time step *t* using Gaussian-sampled noise *ϵ*_*g*_∼ 𝒩 (0, 0.6). We chose this noise source (as well as the photoreceptor noise) such that the model’s spike train variability matched that of the RGCs across repeated stimulus presentations (Extended Data Fig. 1).

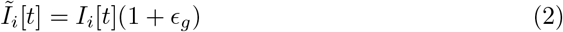

Step 4, RGC transformation: we modelled all RGC spiking units using the normalized and discretized leaky integrate-and-fire model [81]. This evolves the model membrane potential *V*_*i*_[*t*] ∈ ℝ of unit *i* at time *t* using the difference equation:

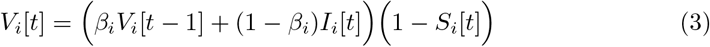

The membrane potential decays by a learnt factor *β*_*i*_ at every simulation step and resets to zero if a spike occurred in the previous simulation step. A spike is emitted if the membrane potential reaches the firing threshold (equal to one).

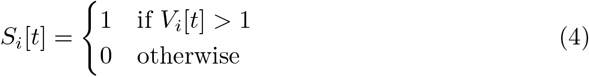

Step 5, stimulus estimation: finally, at every simulation step *t*, a span of *T*_*D*_ = 2 steps of proceeding spike activity, plus a bias *b*^proj^∈ ℝ, was linearly mapped using projection weights 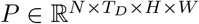 to generate an estimation of the stimulus movie frame 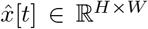 (either of the present or the future, with 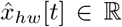 denoting the projected pixel value). This linear mapping is not envisaged as a part of the retinal network, rather it approximates a posited evolutionarily-enforced constraint that causes the network to use representations that can be linearly decoded to provide an estimate of the present or future input [92].

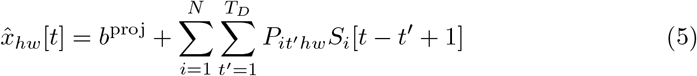

#### Bounding membrane time constants

We bound the *β*_*i*_ value of every spiking unit to an appropriate range to ensure that the network employed biologically-realistic membrane potentials [96].

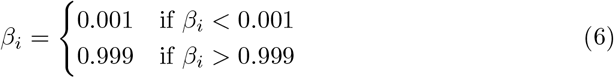

#### Weight initialization

The initial feedforward *W*, recurrent *W* ^rec^ and projection *P* weights were all sampled from a uniform distribution *U* (−*k, k*), with 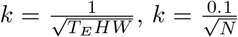 and 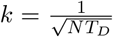 for the feedforward, recurrent and projection weights, respectively. All initial membrane time constants were set to 20ms (*i*.*e. β*_*i*_ ≈ 0.81). All initial bias terms were set to zero.

#### Loss function

The model was trained, by minimizing loss ℒ_total_, to generate a single-frame projection of the natural movie stimulus at every simulation step *t* (corresponding to loss ℒ_projection_) under a metabolic-like constraint (corresponding to loss ℒ_metabolic_), weighted by hyperparameter *λ* = 10^*−*2.5^ (which by qualitative inspection we determined to produce the most retina-like receptive fields across all models).

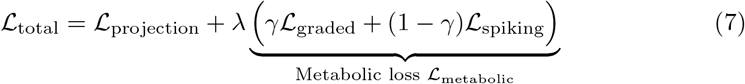

The projection loss ℒ_projection_ is the mean squared error (MSE) between every pixel in the generated single-frame projection 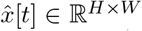 and the corresponding natural stimulus target frame 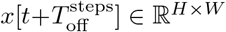 where 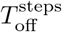 specifies by deviation in time-steps if the frame is of the present (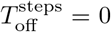) or of the future 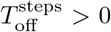 This was computed as the average over all batch samples *B* (which we omit from the notation for brevity), simulation steps *T* (starting from time step *t*_0_ onwards to allow the unit membrane potentials to sufficiently depolarize, *i*.*e*., to permit the network to *warm up*), and spatial height dimension *H* and width dimension *W*, both cropped by *c* = 3 pixels on all sides.

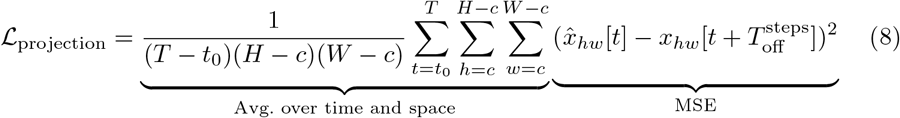

The metabolic-like loss ℒ_metabolic_ is an approximation of the biological energy use of the network. We took this to be a proxy of the energy expenditure at every synapse, calculated as the absolute input value to a synaptic weight, multiplied by its respective absolute value. We weighted the metabolic loss of the graded pre-RGC-like units ℒ_graded_ and the spiking RGC-like units ℒ_spiking_ separately using hyperparameter *γ* = 0.3, as spiking neurons consume more energy than graded neurons [97, 98].

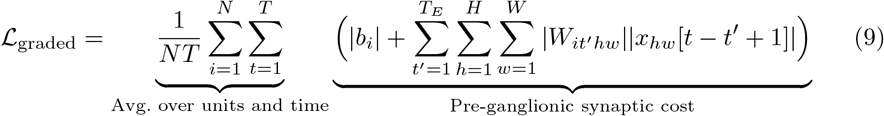

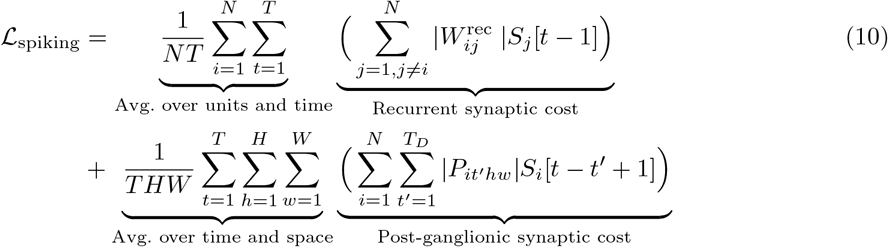

#### Surrogate gradient descent

All spiking retinal models were optimized using a modification of the backpropagation algorithm [99], called surrogate gradient descent [100]. To permit the flow of gradient, the gradient of the non-differentiable Heaviside step function was replaced with a well-behaved function. We adopted the fast sigmoid function, which has been shown to work well in practice [101, 102].

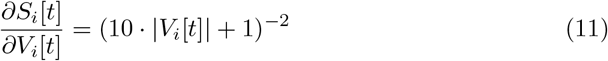

All training was performed using the Adam optimizer [103] (with default parameters) over 100 epochs, with an initial learning rate of 10^*−*4^ (decayed to 10^*−*5^ after 50 epochs) and batch size of 1024. Model weights were saved during training whenever a new minimum training loss was obtained.

### Spike analysis

#### Filtering for active units

For all reported results we only included units that were active and responded to our held-out test set of natural clips. This resulted in 113*/*400 units being dropped from the analysis. All results that report a population percentage are with respect to the number of active units (287).

#### Firing rate

We calculated the firing rate of a neuron (and model unit), by counting the number of spikes over a given stimulus presentation and dividing this by the stimulus duration, to obtain a firing rate.

#### Coefficient of variation

We computed the coefficient of variation of the interspike interval CV(ISI), to quantify the variability of the timing of spikes for a given neuron (and model unit). This is defined as the ratio between the standard deviation and mean of the interspike intervals [26], where a *CV* (*ISI*) *<* 1 corresponds to a regular firing pattern and a *CV* (*ISI*) *>* 1 corresponds to an irregular firing pattern. We excluded CV(ISI) values from the analysis if there were less than three spikes for a given stimulus presentation [104].

#### Fano factor

We reported the Fano factor for each neuron (and model unit) over each stimulus presentation, to estimate the spike count variability over the stimulus-repeat trials. The Fano factor is defined as the variance of the spike count divided by the mean spike count over stimulus-repeat trials.

#### Image decoding from single cell responses

We adopted the experimental paradigm of [21], and obtained the activity of an OFF-type model unit in response to a 150ms-flash of a gray-scale image of a swimming salamander larva. Similar to the experimental paradigm, we convolved the model unit along the x- and y-direction (using a stride of 2 pixels) to obtain the model unit spike activity at different spatial locations. The image was then reconstructed - with every pixel denoting a unique spatial location - using either the number of spikes or the relative timing of the first spikes.

#### Image decoding from population responses to noise-corrupted inputs

We performed all analyses using a selected set of 300 natural photographs from the McGill Calibrated Color Image Database [105]. These images were converted to grayscale as for the model training data, and normalized by subtracting the mean. The spatial dimensions were rescaled to 64×64 pixels using bicubic interpolation, with 240 images used for training and cross-validation and the remaining 60 images used for testing. For every experiment, we appended different sampled Gaussian noise *ϵ* ∼ 𝒩 (0, *σ*_*s*_) to every image pixel, and varied the distribution standard deviation, *σ*_*s*_, from 0 to 2.0 in 0.25 increments between the different experiments. We passed the set of noise-corrupted images into the model (by convolving along the x- and y-direction using a stride of 4 pixels) and transformed the resulting spike outputs into a rate code (*i*.*e*. the number of spikes of a unit) and a relative spike latency code (*i*.*e*. the time of spikes minus the mean spike time across all units to an image). We then fitted a separate linear readout model from each of these respective spike codes, to reconstruct the original images. We penalized the readout weights using an L1 penalty, with the optimal penalty weighting found using five-fold cross-validation. Lastly, we fitted each model using its optimal regularization penalty on all of the training data and reported the mean pixel correlation coefficient of the reconstructed images on the held-out test set.

#### Classifying units with an omitted-stimulus response

RGCs exhibit OSRs at the end of a flash sequence around the expected time of the next flash. These OSRs have been characterized to be stronger than the responses during the flash sequence [22]. Prior studies classified OSRs via qualitative inspection [22, 23]. We developed a set of conditions based on previous descriptions of OSRs and classified units as exhibiting OSRs if they satisfied these conditions. For each unit, we measured the maximum response *R*_*OM*_ during the 80ms duration following the termination of the flash sequence; the maximum response *R*_*F L*_ during the flash sequence; and the baseline response *R*_*BA*_ which we took to be the maximum value during the constant 150ms light flash preceding the flash sequence. The unit was taken to exhibit OSRs if the following conditions were all met: the response directly following the flash sequence was 5% larger than the responses during the flash sequence (*R*_*OM*_ */R*_*F L*_ *>* 1.05); the response directly following the flash sequence was 20% larger than the maximum response to the non-flashing sequence (*R*_*OM*_ */R*_*BA*_ *>* 1.20); and the response directly following the flash sequence was substantial (*R*_*OM*_ *>* 48 spike/s, threshold chosen via qualitative inspection).

### Receptive field analysis

#### Spike-triggered average

We estimated the spatiotemporal receptive field of every unit using a spike-triggered average, using responses to 200 white-noise input clips of 100-frame duration. Clips were generated by sampling each pixel from a Gaussian distribution with a standard deviation of *σ* = 10. We estimated the RF temporal profile by averaging over the spatial pixel values for every frame in time [106].

#### Fitting 2D Gaussians

We fitted a two-dimensional Gaussian function [35, 41] to the spatial RF using gradient descent, with the function defined as:

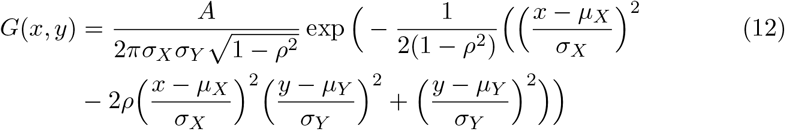

with (*x, y*) denoting the spatial position in pixel space; *A* the amplitude; (*µ*_*X*_, *µ*_*Y*_) the RF centre position; *σ*_*X*_ and *σ*_*Y*_ the standard deviation along the x- and y-axis, respectively, and *ρ* the RF orientation. For each RF, the parameters of the two-dimensional Gaussian were fitted by minimizing the mean squared error between the two-dimensional Gaussian and the RF. For the RF analyses, we only included units (173*/*287) that had a good fit (those with a correlation coefficient *>* 0.7 and those with standard deviation *σ*_*X*_ and *σ*_*Y*_ *>* 0.5 pixels [92]).

#### Classifying unit types

Similarly to classifying the different retinal cell types [35], we classified the functional type of every model unit that had a valid 2D Gaussian fit. For every unit with a sufficient fit, we obtained the spatial RF with the largest power in time, and projected its RF diameter (calculated as 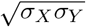) against the first principal component of its temporal profile. We classified the different functional unit types, by grouping the units into four distinct groups using k-means clustering with four fixed centroids.

### Motion tuning

#### Virtual physiology

We measured the model unit response properties to full-field sinusoidal gratings (of amplitude one) of varying orientation (0° to 360° in 5° increments), spatial frequency (10 evenly spaced values between 0.01 to 0.2 cycles per pixel) and temporal frequency (1, 2, 4 and 8Hz). We presented each grating to each unit for 3s and computed the resulting PSTH by averaging over 8 repeats and convolving with a Gaussian kernel (*σ* = 37ms). Next, we computed the mean firing rate over time (ignoring the first 41ms for network warmup, as done in training), resulting in a 3D mean-firing tuning space for each unit (with the preferred stimulus eliciting the highest mean response).

#### Orientation and direction selectivity

We measured the orientation selectivity index (OSI) and direction selectivity index (DSI) for each unit, which respectively quantify the tuning preference for motion across a particular orientation and direction:

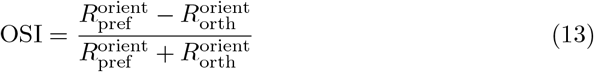

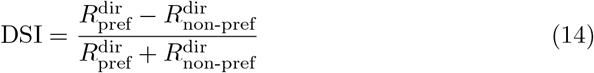

With 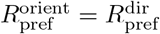 being a unit’s response to a stimulus moving in the preferred direction, and 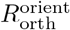 and 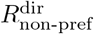 being the responses to the orthogonal orientation and opposite direction, respectively (all using a unit’s preferred spatial and temporal frequency). For both metrics, a value close to zero corresponds to weaker tuning, and a value close to unity corresponds to stronger tuning.

#### Counting Y-type-like units

We presented full-field sinusoidal gratings with a spatial frequency of 0.3 cycles per pixel, at each unit’s preferred orientation. We then measured each unit’s firing rate for the stationary grating *R*_*SG*_ and the grating moving at each unit’s preferred temporal frequency *R*_*MG*_. We considered a unit to be Y-type-like if the following constraints were satisfied: the moving grating response was larger than the stationary grating response (*R*_*MG*_ *> R*_*SG*_); and the stationary and moving grating responses were respectively below (*R*_*SG*_ *<* 24 spikes/s) and above (*R*_*MG*_ *>* 24 spikes/s) a firing threshold (chosen via qualitative inspection) to potentially avoid counting units as Y-type-like due to noise fluctuations.

#### Anticipation code

We implemented the experimental protocol of [24] as closely as possible, where the authors projected a dark bar on a white background moving 6.7*µm* across the retina every 15ms, or where they flashed the bar directly over a unit’s RF for 15ms. We replicated this experiment for the model by projecting a dark bar (pixel value of 1) on a white background (pixel value of−1), moving one pixel every 150ms (36 frames). We calculated this to match the movement speed of the bar in the experiment of [24], assuming an RGC RF diameter of 300*µm* [107] and a model unit RF diameter of 5 pixels, which corresponds to moving the bar by≈0.11 pixel every 4 frames (16.66ms) (or one pixel every 36 frames).

#### Predicting neural responses from model activity

We used publicly available and spike-sorted recordings of RGCs of different animal species in response to natural images and movies. For all datasets, we used 80% of the data for training and the remaining 20% for testing (with splits separated by unique clips or images). As the datasets were recorded at different sampling rates, we resampled all data to be 240Hz (the same sampling rate of the natural movie stimulus with which the spiking retinal model was trained) by repeating frames at particular points in time. We also normalized all natural stimuli (clips and images), by subtracting the mean and dividing by the standard deviation of the training set. We averaged the recorded spike responses over trials and convolved the resulting PSTH with a Gaussian kernel (using *σ* = 38ms, 9 time-steps) to obtain a smoothed spike firing probability for every neuron [37].

#### Macaque image and movie dataset

We used recordings from ON- and OFF-parasol cells in the peripheral retina of macaque monkeys in response to natural images and movies (made available by [36]). This consisted of 16 cells recorded in response to 48 250ms-flashed natural images over 5 trials, and 24 cells recorded in response to 7 6s-long natural movies over 4 trials, with the movies consisting of the natural images spatially shifted in time using eye movement trajectories of human observers [108]. All stimuli were projected at a frame rate of 60Hz onto the receptive field center of each neuron using a circular aperture.

#### Marmoset movie dataset

We used recordings from ON- and OFF-midget and ON- and OFF-parasol cells in the retina of marmoset monkeys in response to natural movies (made available by [37]). This includes 423 cells recorded in response to 22 1s-long natural movies over 30 trials, with the movies consisting of the natural images spatially shifted in time using eye movement trajectories of awake, head-fixed marmoset monkeys. All stimuli were projected at a frame rate of 85Hz. To keep the fitting procedure computationally manageable, we only fitted a subset of the most responsive neurons, as measured by the maximum correlation coefficient (see Section 3), resulting in 164 neurons (using *CC*_max_ ≥ 0.93).

#### Mouse movie dataset

We used recordings from ON- and OFF-midget and ON- and OFF-parasol cells in the mouse retina in response to natural movies (made available by [37]). These 1791 cells were recorded in response to 22 1s-long natural movies over 30 trials, with the movies consisting of the natural images spatially shifted in time using the horizontal gaze component of freely-moving mice. All stimuli were projected at a frame rate of 75Hz. Again, to keep the fitting procedure computationally manageable, we only fitted a subset of the most responsive neurons, as measured by the maximum correlation coefficient (see Section 3), resulting in 351 neurons (using *CC*_max_ ≥ 0.93).

#### Salamander image

We used recordings of various ON- and OFF-type cells in the salamander retina in response to natural images (made available by [35]). This includes 215 cells recorded in response to 300 natural images flashed for 200ms each over 12 trials. There was an 800ms inter-stimulus interval between image presentations, with spike responses to each image recorded between image onset and 100ms after image offset. The images came from the Van Hateren database [109] over 12 trials. All stimuli were projected at a frame rate of 30Hz.

#### Obtaining latent model activity

We trained a set of 14 retinal models, one for every temporal offset *T*_off_ into the future (*T*_off_∈ { 0, 8, 16, 24, 32, 40, 48, 56, 64, 72, 80, 96, 112, 128} ms, where *T*_off_ = 0 corresponds to the present). We convolved each retinal model (using a stride of 4 pixels) over the spatial dimensions of each dataset, to obtain the unit spike activity 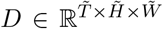 (where the number of simulation steps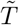, and spatial height 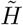 and width 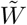 dimensions varied between the different datasets). This was repeated 8 times, and averaged over repeats, to obtain a new activity dataset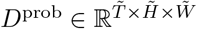, containing the spike probability of every unit over space and time. We further compressed this dataset to a latent activity dataset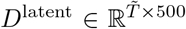, using the first *N*_PC_ = 500 principal components over space. This technique is routinely used [73], as it reduces overfitting [73] and keeps the fitting procedure computationally tractable [110].

#### Fitting procedure

For every retinal model, we fitted a linear-nonlinear readout, to predict the response 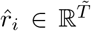 of every neuron *i* in every dataset from the latent model activity. During project development, using only the training set, we explored various hyperparameters relating to the readout model and fitting procedure for each dataset and employed those hyperparameters that we found to work the best.

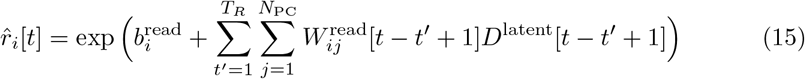

with readout weights 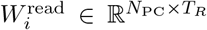 using a *T*_*R*_ = 5 temporal frame span equivalent to≈ 21ms (except for the mouse movie dataset where we used a span of 46ms). Each readout was jointly fitted across all 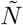 neurons of a dataset, by minimizing the negative Poisson log-likelihood of the readout predictions 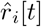 and the neuron’s responses *r*_*i*_[*t*], and an L1 penalty on the readout weights (weighted by hyperparameter *λ*^read^) to avoid overfitting.

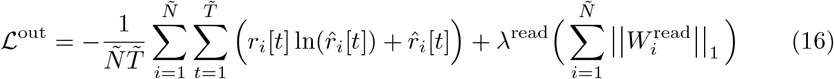

We trained each readout using the Adam optimizer (with default parameters) using a learning rate of 10^*−*4^ over 2000 epochs with a batch size of 8 subsamples of at most 600ms duration (except the salamander dataset where we used a batch size of 32). We found the optimal regularization hyperparameter *λ*^read^ using five-fold cross-validation on the training dataset (over a set of five different log-spaced values, which we varied between the different datasets). Lastly, we also explored different spatial scales of the input stimuli, to account for differences in spatial scale between the model units and neurons [71, 111]. To determine the optimal scale for each dataset, we ran all fits using three different spatial scales (× 0.66, × 1, and× 1.5) using the retinal model optimized for efficient compression of the present (Extended Data Table. 1). We then chose the spatial scale which performed the best on each test dataset to fit the remaining models.

#### Normalized and maximum correlation coefficient

We used the normalized correlation coefficient CC_*norm*_ to quantify the neural fits, expressed as the Pearson correlation coefficient CC between the model’s prediction and target neural response, normalized by dividing by the maximum obtainable correlation coefficient CC_*max*_ to take into account neural noise [72, 112]. A value of zero indicates no correlation between the predicted and target neural response, whereas a value of one indicates the best possible score.

## Acknowledgments

Luke Taylor was supported by the Clarendon Fund of the University of Oxford. Andrew King and Nicol Harper were supported by the Wellcome Trust (WT108369/Z/2015/Z). Friedemann Zenke was supported by Swiss National Science Foundation (grant no. PCEFP3 202981) and the Novartis Research Foundation.

## Data availability

The natural movie training data can be downloaded from https://figshare.com/articles/dataset/Natural_movies/24265498. Pre-trained spiking models can be found at https://github.com/webstorms/RetinalModel. We did not perform any of the retinal recordings. Inspect the data availability sections in the relevant papers cited in this manuscript to access the retinal data. Experimental data in Fig. 4d was extracted using an online tool available at https://apps.automeris.io/wpd/.

## Code availability

The code of the spiking retinal model and the code for reproducing the results can be found at https://github.com/webstorms/RetinalModel.

## Extended data

**Extended Data Table. 1:** Prediction accuracy across different scalings. Prediction score using the encoding retinal model across the different datasets at different spatial scales. Bold values indicate the best performing scale, which was chosen to fit the remaining predictive retinal models in each respective dataset.

| Dataset | Scale | CC<sub>norm</sub> |
| --- | --- | --- |
| Macaque movie | × 0.66 | 0.40 |
|  | × **1.0** | **0.42** |
|  | × 1.5 | 0.39 |
| Macaque image | × 0.66 | 0.60 |
|  | × **1.0** | **0.70** |
|  | × 1.5 | 0.52 |
| Marmoset movie | × 0.66 | 0.24 |
|  | × 1.0 | 0.26 |
|  | × **1.5** | **0.27** |
| Salamander image | × 0.66 | 0.48 |
|  | × **1.0** | **0.52** |
|  | × 1.5 | 0.52 |
| Mouse movie | × 0.66 | 0.13 |
|  | × 1.0 | 0.16 |
|  | × **1.5** | **0.17** |

**Extended Data Fig. 1:**
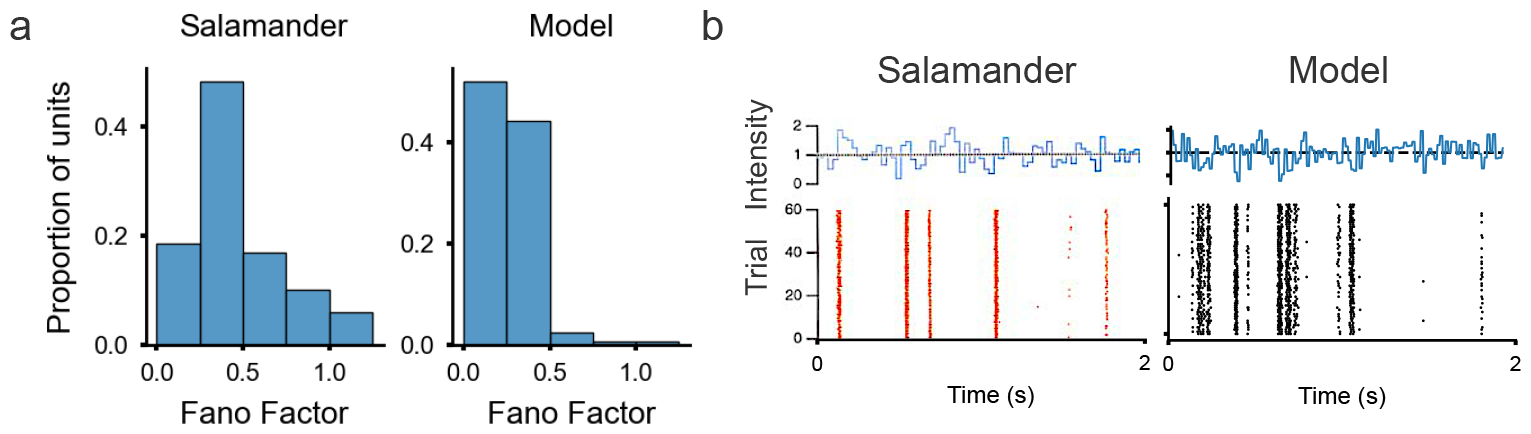
Model response variability. **a**. Histogram of Fano factors in the salamander retina (left) (data from [35]) and the model (right) in response to the same natural images. **b**. Spike raster over 60 trials in response to a uniform full-field illumination of randomly-varying intensity in the salamander retina [3] (left) and the model (right).

**Extended Data Fig. 2:**
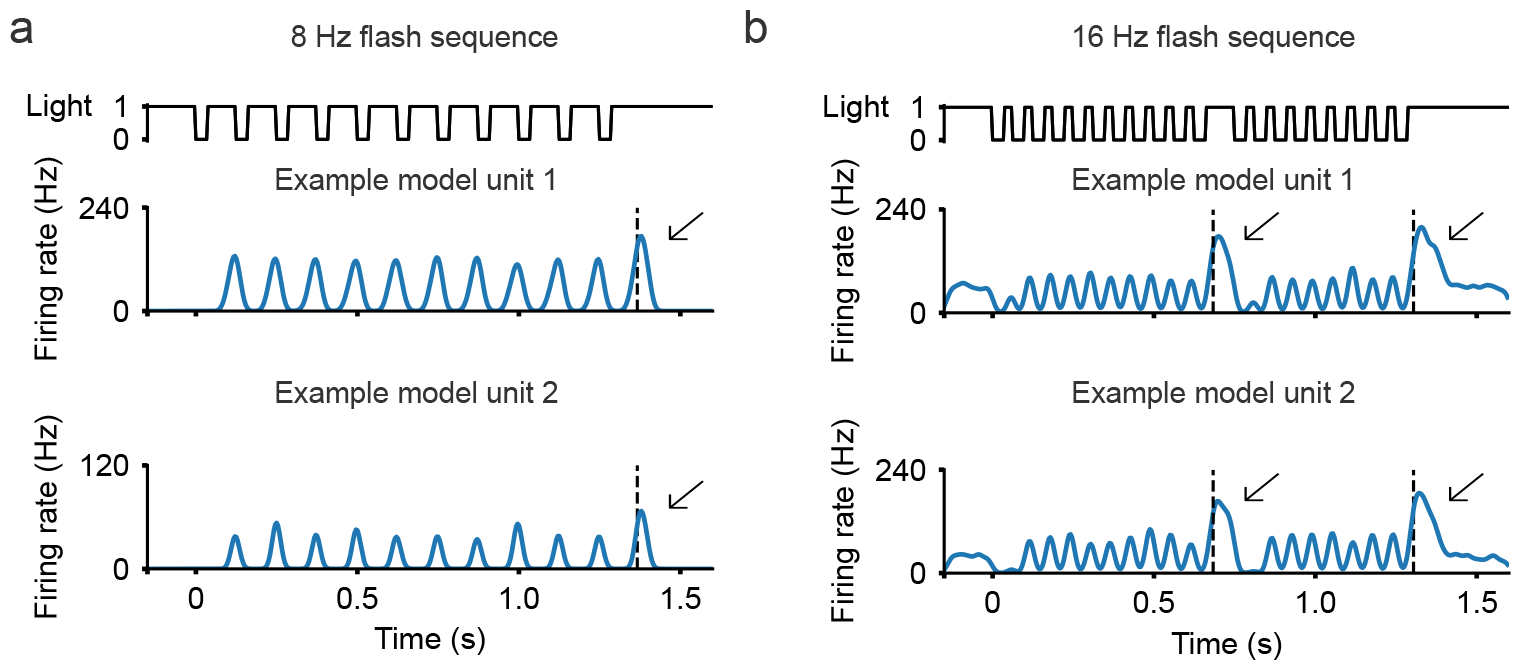
Omission stimulus responses. **a**. Example model unit responses to a sequence flashing at 8 Hz. **b**. Example model unit responses to a sequence flashing at 16 Hz with a flash omission in the middle of the sequence. The dashed line marks the time when the flash sequence was omitted, with the arrow indicating the omitted-stimulus response.

## Notes

https://figshare.com/articles/dataset/Natural_movies/24265498

### Competing Interest Statement

The authors have declared no competing interest.

## References

[1] Hartline, H. K. The response of single optic nerve fibers of the vertebrate eye to illumination of the retina. American Journal of Physiology-Legacy Content 121, 400–415 (1938).

[2] Kuffler, S. W. Discharge patterns and functional organization of mammalian retina. Journal of neurophysiology 16, 37–68 (1953).

[3] Meister, M. & Berry, M. J. The neural code of the retina. neuron 22, 435–450 (1999).

[4] Ocko, S., Lindsey, J., Ganguli, S. & Deny, S. The emergence of multiple retinal cell types through efficient coding of natural movies. Advances in Neural Information Processing Systems 31 (2018).

[5] Jun, N. Y., Field, G. D. & Pearson, J. Scene statistics and noise determine the relative arrangement of receptive field mosaics. Proceedings of the National Academy of Sciences 118, e2105115118 (2021).

[6] Jun, N. Y., Field, G. & Pearson, J. Efficient coding, channel capacity, and the emergence of retinal mosaics. Advances in neural information processing systems 35, 32311–32324 (2022).

[7] Karklin, Y. & Simoncelli, E. Efficient coding of natural images with a population of noisy linear-nonlinear neurons. Advances in neural information processing systems 24 (2011).

[8] Roy, S., Jun, N. Y., Davis, E. L., Pearson, J. & Field, G. D. Inter-mosaic coordination of retinal receptive fields. Nature 592, 409–413 (2021).

[9] Zhaoping, L. Theoretical understanding of the early visual processes by data compression and data selection. Network: computation in neural systems 17, 301–334 (2006).

[10] Barlow, H. B. et al. Possible principles underlying the transformation of sensory messages. Sensory communication 1, 217–233 (1961).

[11] Palmer, S. E., Marre, O., Berry, M. J. & Bialek, W. Predictive information in a sensory population. Proceedings of the National Academy of Sciences 112, 6908–6913 (2015).

[12] Salisbury, J. M. & Palmer, S. E. Optimal prediction in the retina and natural motion statistics. Journal of Statistical Physics 162, 1309–1323 (2016).

[13] Liu, B., Hong, A., Rieke, F. & Manookin, M. B. Predictive encoding of motion begins in the primate retina. Nature neuroscience 24, 1280–1291 (2021).

[14] Berry, M. J. & Schwartz, G. The retina as embodying predictions about the visual world. Predictions in the brain: Using our past to generate a future 295 (2011).

[15] Singer, Y., Taylor, L., Willmore, B. D., King, A. J. & Harper, N. S. Hierarchical temporal prediction captures motion processing along the visual pathway. Elife 12, e52599 (2023).

[16] Keller, G. B. & Mrsic-Flogel, T. D. Predictive processing: a canonical cortical computation. Neuron 100, 424–435 (2018).

[17] Bialek, W., Nemenman, I. & Tishby, N. Predictability, complexity, and learning. Neural computation 13, 2409–2463 (2001).

[18] Dayan, P. & Abbott, L. F. Theoretical neuroscience: computational and mathematical modeling of neural systems (Computational Neuroscience Series, 2001).

[19] Levenstein, D. et al. On the role of theory and modeling in neuroscience. arXiv preprint arXiv:2003.13825 (2020).

[20] Gollisch, T. & Meister, M. Eye smarter than scientists believed: neural computations in circuits of the retina. Neuron 65, 150–164 (2010).

[21] Gollisch, T. & Meister, M. Rapid neural coding in the retina with relative spike latencies. science 319, 1108–1111 (2008).

[22] Schwartz, G., Harris, R., Shrom, D. & Berry, M. J. Detection and prediction of periodic patterns by the retina. Nature neuroscience 10, 552–554 (2007).

[23] Schwartz, G. & Berry 2nd, M. J. Sophisticated temporal pattern recognition in retinal ganglion cells. Journal of neurophysiology 99, 1787–1798 (2008).

[24] Berry, M. J., Brivanlou, I. H., Jordan, T. A. & Meister, M. Anticipation of moving stimuli by the retina. Nature 398, 334–338 (1999).

[25] Ölveczky, B. P., Baccus, S. A. & Meister, M. Segregation of object and background motion in the retina. Nature 423, 401–408 (2003).

[26] Gerstner, W., Kistler, W. M., Naud, R. & Paninski, L. Neuronal dynamics: From single neurons to networks and models of cognition (Cambridge University Press, 2014).

[27] Baccus, S. A. & Meister, M. Fast and slow contrast adaptation in retinal circuitry. Neuron 36, 909–919 (2002).

[28] Pillow, J. W. et al. Spatio-temporal correlations and visual signalling in a complete neuronal population. Nature 454, 995–999 (2008).

[29] Vaney, D. I. Patterns of neuronal coupling in the retina. Progress in retinal and eye research 13, 301–355 (1994).

[30] Vlasiuk, A. & Asari, H. Feedback from retinal ganglion cells to the inner retina. PLoS One 16, e0254611 (2021).

[31] Baylor, D., Matthews, G. & Yau, K. Two components of electrical dark noise in toad retinal rod outer segments. The Journal of physiology 309, 591–621 (1980).

[32] van Rossum, M. C., O’Brien, B. J. & Smith, R. G. Effects of noise on the spike timing precision of retinal ganglion cells. Journal of neurophysiology 89, 2406–2419 (2003).

[33] Berry, M. J., Warland, D. K. & Meister, M. The structure and precision of retinal spike trains. Proceedings of the National Academy of Sciences 94, 5411–5416 (1997).

[34] Berry, M. & Meister, M. Refractoriness and neural precision. Advances in neural information processing systems 10 (1997).

[35] Liu, J. K., Karamanlis, D. & Gollisch, T. Simple model for encoding natural images by retinal ganglion cells with nonlinear spatial integration. PLoS Computational Biology 18, e1009925 (2022).

[36] Freedland, J. & Rieke, F. Systematic reduction of the dimensionality of natural scenes allows accurate predictions of retinal ganglion cell spike outputs. Proceedings of the National Academy of Sciences 119, e2121744119 (2022).

[37] Karamanlis, D. et al. Natural stimuli drive concerted nonlinear responses in populations of retinal ganglion cells. bioRxiv 2023–01 (2023).

[38] Dacey, D. M. The mosaic of midget ganglion cells in the human retina. Journal of Neuroscience 13, 5334–5355 (1993).

[39] Dacey, D. 20 origins of perception: Retinal ganglion cell diversity and the creation of parallel visual pathways (2004).

[40] Soto, F. et al. Efficient coding by midget and parasol ganglion cells in the human retina. Neuron 107, 656–666 (2020).

[41] Kling, A. et al. Functional organization of midget and parasol ganglion cells in the human retina. BioRxiv 2020–08 (2020).

[42] Lee, B. B. Receptive field structure in the primate retina. Vision research 36, 631–644 (1996).

[43] Field, G. D. et al. Spatial properties and functional organization of small bistratified ganglion cells in primate retina. Journal of Neuroscience 27, 13261–13272 (2007).

[44] Devries, S. H. & Baylor, D. A. Mosaic arrangement of ganglion cell receptive fields in rabbit retina. Journal of neurophysiology 78, 2048–2060 (1997).

[45] Wässle, H., Peichl, L. & Boycott, B. B. Morphology and topography of on-and off-alpha cells in the cat retina. Proceedings of the Royal Society of London. Series B. Biological Sciences 212, 157–175 (1981).

[46] Field, G. D. & Chichilnisky, E. Information processing in the primate retina: circuitry and coding. Annu. Rev. Neurosci. 30, 1–30 (2007).

[47] Dacey, D. M. Physiology, morphology and spatial densities of identified ganglion cell types in primate retina, 12–34 (Wiley Online Library, 2007).

[48] Ratliff, C. P., Borghuis, B. G., Kao, Y.-H., Sterling, P. & Balasubramanian, V. Retina is structured to process an excess of darkness in natural scenes. Proceedings of the National Academy of Sciences 107, 17368–17373 (2010).

[49] O’Brien, B. J., Isayama, T., Richardson, R. & Berson, D. M. Intrinsic physiological properties of cat retinal ganglion cells. The Journal of physiology 538, 787–802 (2002).

[50] Sabbah, S. et al. A retinal code for motion along the gravitational and body axes. Nature 546, 492–497 (2017).

[51] Oyster, C. W. & Barlow, H. B. Direction-selective units in rabbit retina: distribution of preferred directions. Science 155, 841–842 (1967).

[52] Kühn, N. K. & Gollisch, T. Joint encoding of object motion and motion direction in the salamander retina. Journal of Neuroscience 36, 12203–12216 (2016).

[53] Maximov, V., Maximova, E. & Maximov, P. Direction selectivity in the goldfish tectum revisited. Annals of the New York Academy of Sciences 1048, 198–205 (2005).

[54] Bowling, D. Light responses of ganglion cells in the retina of the turtle. The Journal of Physiology 299, 173–196 (1980).

[55] Nath, A. & Schwartz, G. W. Cardinal orientation selectivity is represented by two distinct ganglion cell types in mouse retina. Journal of Neuroscience 36, 3208–3221 (2016).

[56] Levick, W. R. Receptive fields and trigger features of ganglion cells in the visual streak of the rabbit’s retina. The Journal of physiology 188, 285 (1967).

[57] Amthor, F. R., Takahashi, E. S. & Oyster, C. W. Morphologies of rabbit retinal ganglion cells with complex receptive fields. Journal of comparative neurology 280, 97–121 (1989).

[58] Rosenberg, A. & Talebi, V. The primate retina contains distinct types of y-like ganglion cells. Journal of Neuroscience 29, 5048–5050 (2009).

[59] Enroth-Cugell, C. & Robson, J. G. The contrast sensitivity of retinal ganglion cells of the cat. The Journal of physiology 187, 517–552 (1966).

[60] Caldwell, J. & Daw, N. New properties of rabbit retinal ganglion cells. The Journal of Physiology 276, 257–276 (1978).

[61] Petrusca, D. et al. Identification and characterization of a y-like primate retinal ganglion cell type. Journal of Neuroscience 27, 11019–11027 (2007).

[62] Lettvin, J. Y., Maturana, H. R., McCulloch, W. S. & Pitts, W. H. What the frog’s eye tells the frog’s brain. Proceedings of the IRE 47, 1940–1951 (1959).

[63] Baccus, S. A., Ö lveczky, B. P., Manu, M. & Meister, M. A retinal circuit that computes object motion. Journal of Neuroscience 28, 6807–6817 (2008).

[64] Dayan, P. & Abbott, L. F. Theoretical neuroscience: computational and mathematical modeling of neural systems (MIT press, 2005).

[65] Stanley, G. B. Reading and writing the neural code. Nature neuroscience 16, 259–263 (2013).

[66] Judge, S. J., Richmond, B. J. & Chu, F. C. Implantation of magnetic search coils for measurement of eye position: an improved method. Vision research (1980).

[67] Martinez-Conde, S., Macknik, S. L. & Hubel, D. H. The role of fixational eye movements in visual perception. Nature reviews neuroscience 5, 229–240 (2004).

[68] Karamanlis, D., Schreyer, H. M. & Gollisch, T. Retinal encoding of natural scenes. Annual Review of Vision Science 8, 171–193 (2022).

[69] Sterling, P. & Laughlin, S. Principles of neural design (MIT press, 2015).

[70] Auksztulewicz, R., Rajendran, V. G., Peng, F., Schnupp, J. W. H. & Harper, N. S. Omission responses in local field potentials in rat auditory cortex. BMC biology 21, 130 (2023).

[71] Cadena, S. A. et al. Deep convolutional models improve predictions of macaque v1 responses to natural images. PLoS computational biology 15, e1006897 (2019).

[72] Schoppe, O., Harper, N. S., Willmore, B. D., King, A. J. & Schnupp, J. W. Measuring the performance of neural models. Frontiers in computational neuroscience 10, 10 (2016).

[73] Schrimpf, M. et al. Brain-score: Which artificial neural network for object recognition is most brain-like? BioRxiv 407007 (2018).

[74] Kerschensteiner, D. Feature detection by retinal ganglion cells. Annual review of vision science 8, 135–169 (2022).

[75] Tanaka, H. et al. From deep learning to mechanistic understanding in neuroscience: the structure of retinal prediction. Advances in neural information processing systems 32 (2019).

[76] Maheswaranathan, N. et al. Interpreting the retinal neural code for natural scenes: From computations to neurons. Neuron 111, 2742–2755 (2023).

[77] Schnapf, J., Kraft, T. & Baylor, D. Spectral sensitivity of human cone photoreceptors. Nature 325, 439–441 (1987).

[78] Donner, K. Temporal vision: measures, mechanisms and meaning. Journal of Experimental Biology 224, jeb222679 (2021).

[79] Shapley, R. M. & Victor, J. D. The effect of contrast on the transfer properties of cat retinal ganglion cells. The Journal of physiology 285, 275–298 (1978).

[80] Kim, K. J. & Rieke, F. Temporal contrast adaptation in the input and output signals of salamander retinal ganglion cells. Journal of Neuroscience 21, 287–299 (2001).

[81] Taylor, L., King, A. J. & Harper, N. S. Addressing the speed-accuracy simulation trade-off for adaptive spiking neurons. arXiv preprint arXiv:2311.11390 (2023).

[82] Schreyer, H. M. & Gollisch, T. Nonlinear spatial integration in retinal bipolar cells shapes the encoding of artificial and natural stimuli. Neuron 109, 1692–1706 (2021).

[83] Dacey, D. M. & Packer, O. S. Colour coding in the primate retina: diverse cell types and cone-specific circuitry. Current opinion in neurobiology 13, 421–427 (2003).

[84] Krizhevsky, A., Sutskever, I. & Hinton, G. E. Imagenet classification with deep convolutional neural networks. Advances in neural information processing systems 25 (2012).

[85] Baden, T., Euler, T. & Berens, P. Understanding the retinal basis of vision across species. Nature Reviews Neuroscience 21, 5–20 (2020).

[86] Perez-Nieves, N. & Goodman, D. Sparse spiking gradient descent. Advances in Neural Information Processing Systems 34, 11795–11808 (2021).

[87] Chichilnisky, E. & Kalmar, R. S. Functional asymmetries in on and off ganglion cells of primate retina. Journal of Neuroscience 22, 2737–2747 (2002).

[88] Shi, Q., Gupta, P., Boukhvalova, A. K., Singer, J. H. & Butts, D. A. Functional characterization of retinal ganglion cells using tailored nonlinear modeling. Scientific reports 9, 8713 (2019).

[89] Parikh, N., Itti, L. & Weiland, J. Saliency-based image processing for retinal prostheses. Journal of neural engineering 7, 016006 (2010).

[90] Feng, C., Dai, S., Zhao, Y. & Liu, S. Edge-preserving image decomposition based on saliency map, 159–163 (IEEE, 2014).

[91] Erickson-Davis, C. & Korzybska, H. What do blind people “see” with retinal prostheses. Observations and qualitative reports of epiretinal implant users (Neuroscience) (2020).

[92] Singer, Y. et al. Sensory cortex is optimized for prediction of future input. Elife 7, e31557 (2018).

[93] Taylor, L. & Nitschke, G. Improving deep learning with generic data augmentation, 1542–1547 (IEEE, 2018).

[94] Baden, T., Esposti, F., Nikolaev, A. & Lagnado, L. Spikes in retinal bipolar cells phase-lock to visual stimuli with millisecond precision. Current Biology 21, 1859–1869 (2011).

[95] Ganczer, A. et al. Transiency of retinal ganglion cell action potential responses determined by psth time constant. Plos one 12, e0183436 (2017).

[96] Perez-Nieves, N., Leung, V. C., Dragotti, P. L. & Goodman, D. F. Neural heterogeneity promotes robust learning. Nature communications 12, 5791 (2021).

[97] Laughlin, S. B., de Ruyter van Steveninck, R. R. & Anderson, J. C. The metabolic cost of neural information. Nature neuroscience 1, 36–41 (1998).

[98] Niven, J. E. & Laughlin, S. B. Energy limitation as a selective pressure on the evolution of sensory systems. Journal of Experimental Biology 211, 1792–1804 (2008).

[99] Rumelhart, D. E., Hinton, G. E. & Williams, R. J. Learning representations by back-propagating errors. nature 323, 533–536 (1986).

[100] Neftci, E. O., Mostafa, H. & Zenke, F. Surrogate gradient learning in spiking neural networks: Bringing the power of gradient-based optimization to spiking neural networks. IEEE Signal Processing Magazine 36, 51–63 (2019).

[101] Zenke, F. & Ganguli, S. Superspike: Supervised learning in multilayer spiking neural networks. Neural computation 30, 1514–1541 (2018).

[102] Zenke, F. & Vogels, T. P. The remarkable robustness of surrogate gradient learning for instilling complex function in spiking neural networks. Neural computation 33, 899–925 (2021).

[103] Kingma, D. P. & Ba, J. Adam: A method for stochastic optimization. arXiv preprint arXiv:1412.6980 (2014).

[104] Rasch, M. J., Schuch, K., Logothetis, N. K. & Maass, W. Statistical comparison of spike responses to natural stimuli in monkey area v1 with simulated responses of a detailed laminar network model for a patch of v1. Journal of neurophysiology 105, 757–778 (2011).

[105] Olmos, A. & Kingdom, F. A. A biologically inspired algorithm for the recovery of shading and reflectance images. Perception 33, 1463–1473 (2004).

[106] Ruda, K., Rudzite, A. M. & Field, G. D. The functional organization of retinal ganglion cell receptive fields across light levels. bioRxiv 2022–09 (2022).

[107] Buldyrev, I. & Taylor, W. R. Inhibitory mechanisms that generate centre and surround properties in on and off brisk-sustained ganglion cells in the rabbit retina. The Journal of physiology 591, 303–325 (2013).

[108] Van Der Linde, I., Rajashekar, U., Bovik, A. C. & Cormack, L. K. Doves: a database of visual eye movements. Spatial vision 22, 161 (2009).

[109] Van Hateren, J. H. & van der Schaaf, A. Independent component filters of natural images compared with simple cells in primary visual cortex. Proceedings of the Royal Society of London. Series B: Biological Sciences 265, 359–366 (1998).

[110] Storrs, K. R., Kietzmann, T. C., Walther, A., Mehrer, J. & Kriegeskorte, N. Diverse deep neural networks all predict human inferior temporal cortex well, after training and fitting. Journal of cognitive neuroscience 33, 2044–2064 (2021).

[111] Mineault, P., Bakhtiari, S., Richards, B. & Pack, C. Your head is there to move you around: Goal-driven models of the primate dorsal pathway. Advances in Neural Information Processing Systems 34, 28757–28771 (2021).

[112] Hsu, A., Borst, A. & Theunissen, F. E. Quantifying variability in neural responses and its application for the validation of model predictions. Network: Computation in Neural Systems 15, 91–109 (2004).

